# A Modular Toolkit For Theophylline-Dependent Synthetic Auxotrophs Via Riboswitch-Gated Essential Genes

**DOI:** 10.1101/2025.11.20.689564

**Authors:** Carlos Gonzalez-Lopez, Adrian Overly, Saahil Singh, Charlie Huang, Gabriel Lopez

**Author notes:** Corresponding authors (CH), (GL). These authors contributed equally as last authors (Charlie Huang, Gabriel Lopez).

## Abstract

Synthetic auxotrophs are a useful means for developing genetically encoded biocontainment systems. Current methods for developing synthetic auxotrophs are complicated or expensive, limiting access and adoption of biocontainment technologies. To address this gap, we developed a simple and modular platform for creating synthetic auxotrophs based on ligand-dependent translational gating of essential genes. By inserting a theophylline-responsive riboswitch upstream of essential genes in *Escherichia coli*, we created strains whose viability depends on the presence of theophylline. We systematically applied this approach to 29 essential genes, obtained ligand-dependent growth phenotypes for 19 targets, and found that 18 of these essential genes yielded stringent live-die synthetic auxotrophs. These strains exhibited robust theophylline dependence with escape frequencies ranging from 1 × 10^−5^ to 1 × 10^−6^ (most below limit of detection). Our modular design allows for the rapid (<1 week), low-cost, and reliable generation of synthetic auxotrophs. This work introduces ligand-dependent translational control as a new mechanism for engineering synthetic auxotrophy and provides an accessible platform that expands the biocontainment toolkit.

## INTRODUCTION

### A synthetic biology safety challenge: the need for biocontainment

As synthetic biology expands into medicine, agriculture, and manufacturing, the need for robust containment of genetically engineered organisms becomes increasingly critical [1, 2]. Biocontainment strategies typically aim to prevent escape of engineered organisms. Such safeguards can help prevent accidental release into the environment, avert potential harm to human and animal health, or protect intellectual property from biopiracy.

### Overview of biocontainment approaches and their limitations

Physical containment strategies, such as laboratory infrastructure and operational protocols, provide an external layer of control but are subject to operational or equipment failure. Genetically encoded biocontainment strategies address this limitation by engineering containment directly into an organism’s genome. These strategies are classified into two primary categories: active and intrinsic/passive [2, 3].

Active biocontainment systems typically use genetic circuits to induce cell death in response to an environmental signal (e.g. “kill switches”) [4–6]. Common implementations include toxin-antitoxin modules or the conditional expression of nucleases that degrade the genome [6–8]. The primary limitations of active systems are their requirement for human intervention (meaning organisms live by default) as well as their reliance on multi-component circuits that can be disabled through loss-of-function mutations. Because the biocontainment system is not essential for viability, there is strong selective pressure to inactivate it.

In contrast, intrinsic biocontainment systems typically exploit engineered dependencies for viability, for example through creation of a dependency on an exogenously supplied molecule or condition. In the absence of this input, the organism cannot grow or survive (i.e. it dies by default). This approach, analogous to a dead-man’s switch, provides a more robust framework for containment because death is the default state and does not require an active killing process to function [1, 2].

A key implementation of intrinsic biocontainment is synthetic auxotrophy. This must be distinguished from natural auxotrophy, which involves metabolic dependencies (e.g. amino acids). Natural auxotrophs are unsuitable for robust biocontainment because the required molecules can often be acquired from the environment through scavenging or cross-feeding from other microbes [9, 10].

This limitation led to the development of synthetic auxotrophy, which we define as an engineered chemical dependency on a molecule that is not a normal part of an organism’s metabolism or for which the dependency is non-metabolic. This has so far been accomplished by imposing chemical control over the expression (e.g., transcription, translational elongation) or function (e.g., post-translational) of essential genes and their products. For robust containment, this dependency should be engineered to a molecule that is absent from natural environments and cannot be acquired through cross-feeding or scavenging [4, 11–15].

To date, synthetic auxotrophy has been implemented through several mechanisms that operate at distinct levels of biological control. These include: Non-Standard Amino Acid (NSAA) Dependence, an approach that operates at the level of translational elongation. The genetic code of an organism is expanded, allowing incorporation of NSAAs within essential genes, typically by reassigning a stop codon. In the absence of the externally supplied NSAA, translation of these essential proteins terminates prematurely, causing cell death. In the presence of NSAA, translation occurs and the NSAA is incorporated into the essential gene product. This method can achieve low escape frequencies, but its utility is limited by the requirement for extensively recoded genomes, a significant technical barrier [11, 12, 16, 17].

SLiDE (Synthetic auxotrophs based on Ligand-Dependent Essential genes) imposes chemical control over post translational essential gene product function. An essential protein is engineered to be conformationally unstable and inactive by default (resulting in lethality), but the addition of a specific, synthetic small molecule binds to the engineered protein, restoring its stability and function. This strategy of chemically reducing essential gene product function is more accessible as it does not require genome recoding, but it necessitates bespoke protein engineering and screening for each new essential gene target [14]. More recent approaches have extended this approach using modular approaches [15, 16, 18].

Mapping synthetic auxotroph biocontainment strategies onto the central dogma reveals that multiple control points have been successfully exploited: transcriptional regulation (conditionally expressed essential genes), translational elongation (NSAA addiction), and post-translational protein function (SLiDE). However, translation initiation, where ribosomes are recruited to mRNA, has not been exploited as a means of developing synthetic auxotrophy [19, 20], with the closest previous work using riboswitches to create conditional hypomorphs for functional genomics rather than biocontainment [21]. That work intentionally preserved sufficient gene expression in the restrictive condition in order to enable survival and phenotype analysis. Additionally, a gap remains for exceptionally easy means of engineering intrinsically contained living systems. These established approaches present a trade-off between containment stringency and the ease and modularity of implementation. Here, we address these gaps by introducing a new class of synthetic auxotrophs based on ligand-gated control of translation initiation.

### Approach: Riboswitch Translational Initiation Control as a New Strategy for Synthetic Auxotrophy

Ligand-responsive riboswitches are a molecular tool for imposing ligand-dependent control over translational initiation of essential gene products. Riboswitches are naturally occurring or engineered RNA regulatory elements that function as conformational switches in response to molecular binding. Typically located in the 5’ untranslated region (UTR) of mRNAs, these structured RNAs directly couple ligand recognition to translation. In the absence of ligand, the riboswitch structure sequesters the ribosome binding site (RBS), preventing translation. Upon ligand binding, the RNA undergoes a conformational change that exposes the RBS, permitting translation to proceed.

Our core hypothesis is that inserting a ligand-responsive riboswitch upstream of an essential gene at its native chromosomal locus will create a synthetic auxotroph whose viability is dependent on the riboswitch’s ligand. While many characterized riboswitches are available, we selected the theophylline-responsive riboswitch (riboA) for this demonstration based on practical considerations: it is well-characterized in synthetic biology applications and it responds to a safe, cheap, available compound at practical working concentrations (0.5-2 mM) [22–26].

Our proposed riboswitch-based translational gating strategy addresses the identified need for a more accessible, modular biocontainment platform. The approach is modular because the same riboswitch cassette functions across different essential gene targets without requiring target-specific engineering. Once the riboswitch-RBS architecture is optimized, it can be deployed to any essential gene by simply changing the flanking homology arms in the PCR primers. There is no need for protein engineering, directed evolution, or empirical screening because the RNA-based mechanism is portable across targets [23]. The regulatory mechanism is based on RNA secondary structure, which is well-understood and computationally tractable. Integration follows standardized lambda Red recombination protocols, and outcomes are consistent across gene targets. Implementation requires only standard lambda Red recombination, a routine technique available in any *E. coli* strain carrying the pKD46 helper plasmid [27, 28]. There is no requirement for genome recoding, specialized reagents, complex multi-step selections, or labor-intensive screening procedures. New synthetic auxotrophs can be generated rapidly at low cost using commodity molecular biology tools, which enables parallel construction and testing of numerous candidate biocontainment strains.

We describe the development and validation of a modular, low-cost, rapid toolkit for creating synthetic auxotrophs using theophylline-dependent riboswitches to impose translationally-gated control over essential gene product expression. In order to create the toolkit, we developed a standardized parts kit that enables one-step generation of synthetic auxotrophs (Fig 1). Using this toolkit, we targeted 29 essential genes in *E. coli* and observed ligand-dependent phenotypes for 19 of them. Of these, 18 essential gene targets resulted in strict live-die behavior in response to theophylline.

**Figure 1:**
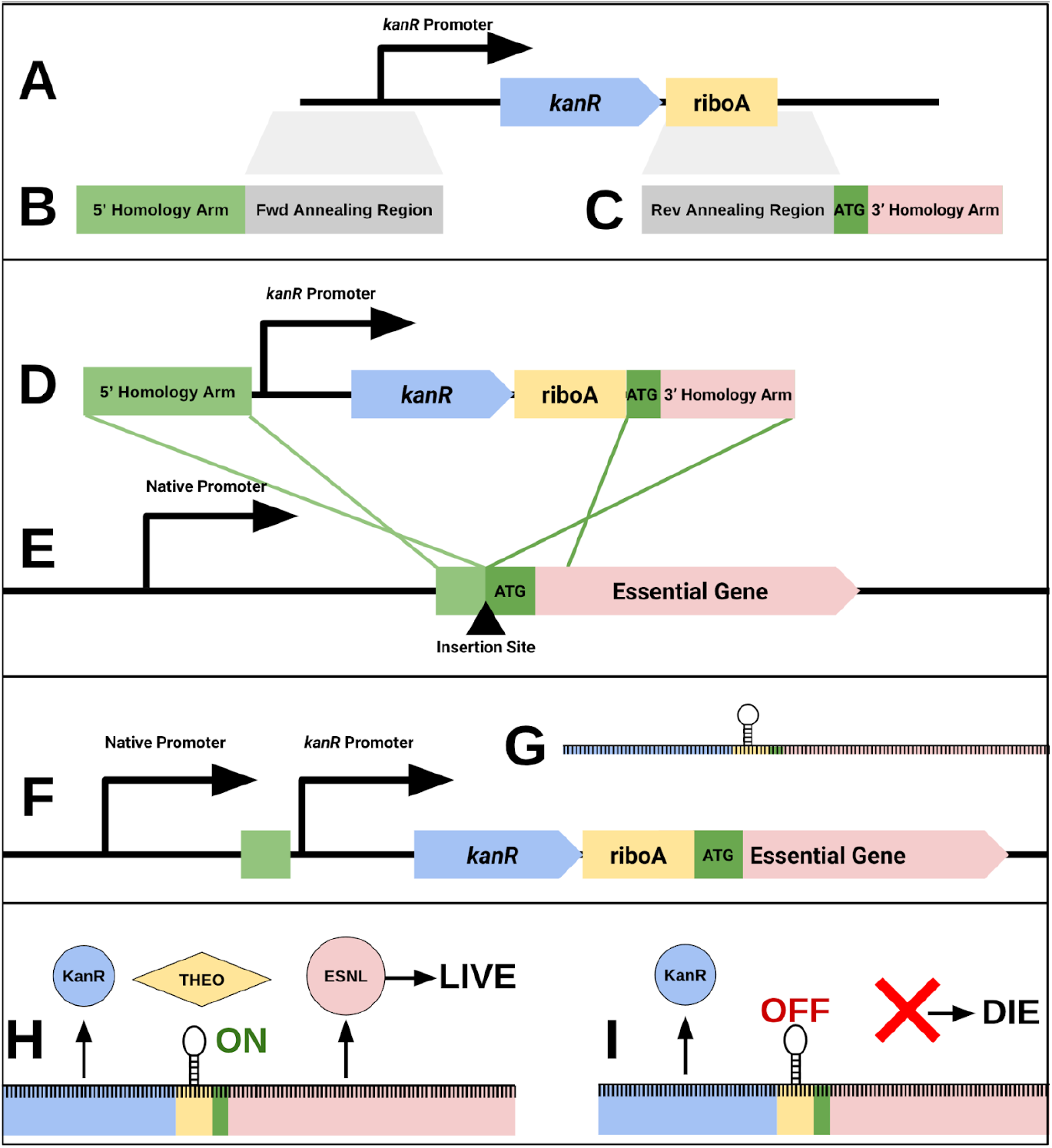
Engineering and Mechanism of Theophylline-Dependent Synthetic Auxotrophy. (A) Template DNA containing a kanamycin resistance marker (kanR) and a theophylline riboswitch (riboA). (B, C) Forward (B) and reverse (C) primers amplify the kanR-riboA cassette. Trapezoids indicate annealing regions. The primers add 5’ and 3’ homology arms targeting the genomic region upstream of an essential gene. (D) The resulting linear PCR product is the kanR-riboA cassette flanked by homology arms. (E) Lambda Red recombination integrates the PCR product into the E. coli chromosome. Homology arms direct insertion immediately upstream of the essential gene’s start codon, and the kanR marker enables antibiotic selection. (F) In the final engineered locus, a constitutive promoter drives transcription of the kanR-riboA-essential gene cassette into a polycistronic-like mRNA (G). (G) The resulting mRNA. kanR (blue) is constitutively translated. The riboA riboswitch (yellow) forms a hairpin, sequestering the RBS of the essential gene (pink) and blocking translation. The start codon is green. (H) Permissive conditions: Theophylline (diamond) binding induces a conformational change (ON), exposing the RBS and enabling translation of the essential protein, which ensures cell viability. (I) Restrictive conditions: Without theophylline, the riboswitch remains closed (OFF), blocking the RBS. Essential gene translation is repressed, leading to cell death.

The toolkit consists of a template plasmid containing a kanamycin resistance marker (*kanR*) and theophylline-responsive riboswitch (riboA). To create a knock-in fragment targeting a specific essential gene, researchers design PCR primers that anneal to the *kanR*-riboA template and include 5’ and 3’ homology arms matching the genomic regions flanking the target gene’s start codon. PCR amplification generates a linear fragment in which the *kanR*-riboA cassette is flanked by gene-specific homology arms. This fragment is integrated immediately upstream of the essential gene’s start codon using lambda Red recombination. Selection on kanamycin plus theophylline yields theophylline-dependent strains in a single step.

We systematically applied this toolkit across 29 essential *E. coli* genes selected to represent diverse cellular functions (Table 1)[29]. This breadth of targets tests whether the riboswitch-gating approach is truly generalizable and illustrates expected success rates for any given attempt. It should be noted that there are a number of different riboswitch variants that could be used in order to tune the gating level, but we decided to standardize on riboA.

**Table 1:**
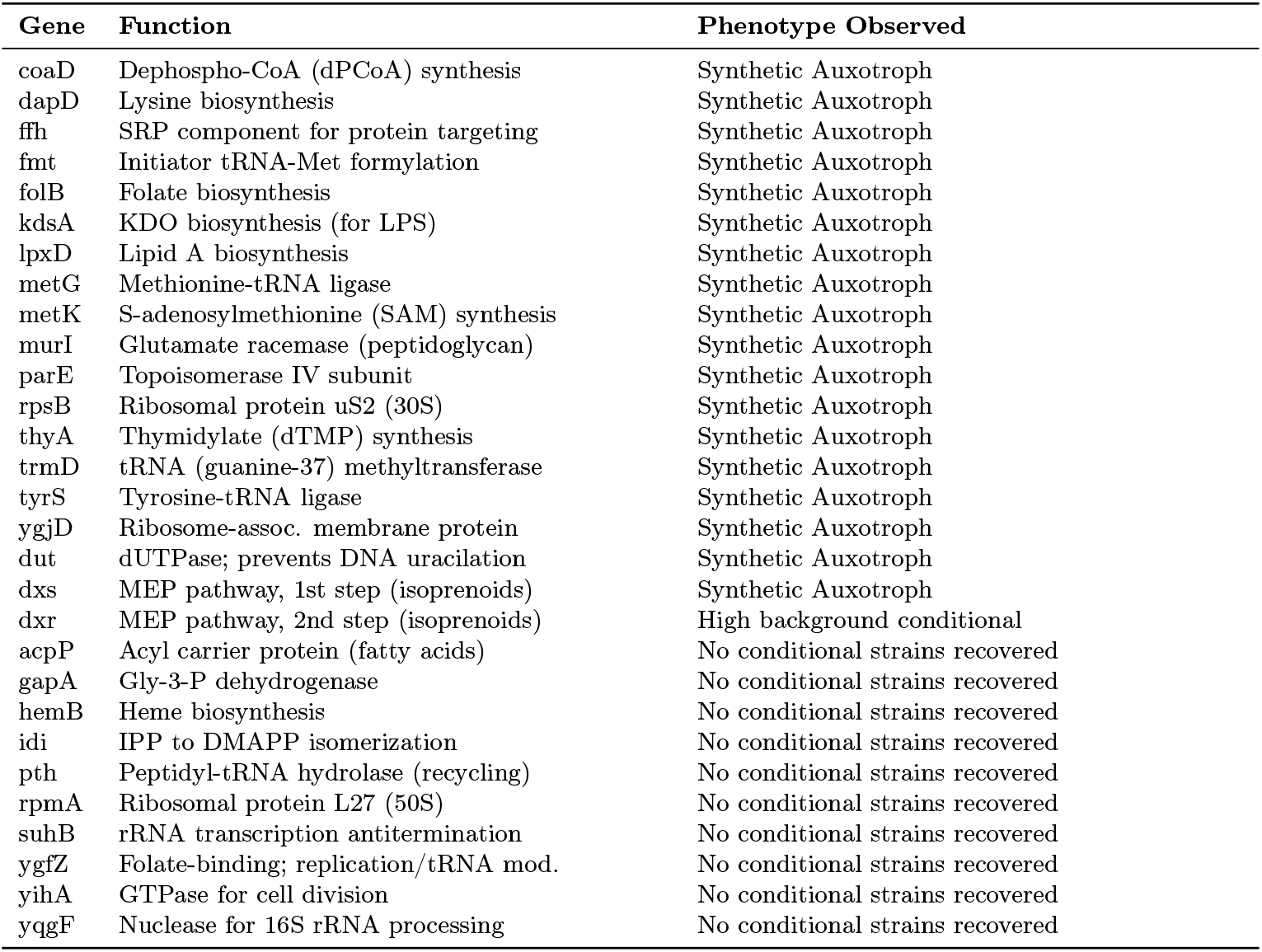
Phenotypic outcomes of riboswitch insertion upstream of essential genes. This table summarizes the results of inserting the kanR-riboA cassette upstream of 29 essential E. coli genes. The targeted genes represent diverse cellular functions, including translation, metabolism, and cell wall biosynthesis. Outcomes are categorized as Theophylline-Dependent Synthetic Auxotroph (viability requires theophylline), Ligand-Modulated Growth (growth is impaired but not eliminated without theophylline), or Non-conditional (insertion attempts did not result in theophylline-dependence). This final category describes targets where either very few viable colonies were recovered, suggesting the intended insertion is lethal, or where numerous viable insertions were obtained, but the resulting strains were non-conditional, indicating that basal gene expression was sufficient to support viability.

From the 29 genes tested, we successfully generated 18 distinct synthetic auxotrophs that demonstrated robust theophylline-dependent viability. These strains formed normal-looking colonies on petri dishes supplemented with theophylline, but failed to grow in its absence, confirming successful translational gating (Fig 2). The recovery rate demonstrates that riboswitch-based translational control is feasible across a substantial fraction of essential genes, with failures attributable to context-dependent factors such as UTR structure, gene expression levels, and protein stability.

**Figure 2:**
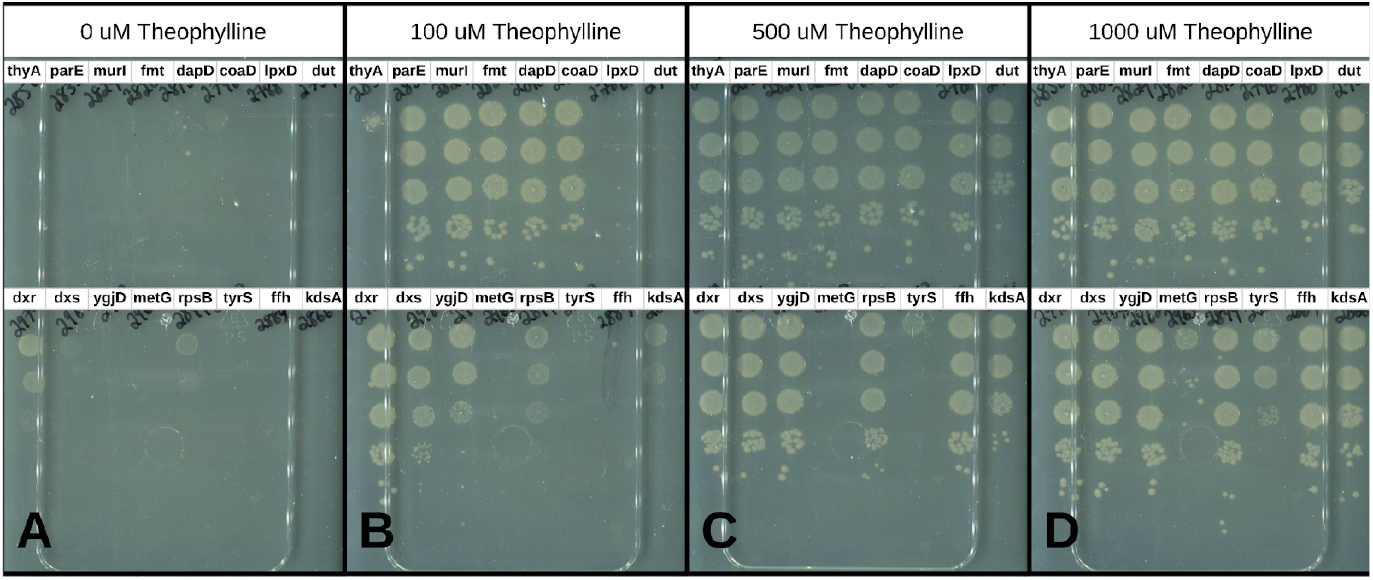
Representative panel of 16 of 19 theophylline-dependent strains grown on restrictive and a titration of permissive condition LB agar petri dishes. This set of petri dishes illustrates viability profiles of a panel of 16 riboA-gated essential genes of which 14 are synthetic auxotrophs. Serial dilutions of each strain were spotted onto petri dishes with various theophylline concentrations. (A) Plates that do not contain theophylline, resulted in no growth for 14/16 strains and background growth for dxr and rpsB. (B) Plates supplemented with 100 uM of theophylline resulted in full to partial rescue of ∼9/16 strains. (C) Plates supplemented with 500 uM of theophylline resulted in rescue of 14/16 strains. (D) Plates supplemented with 1000 uM of theophylline resulted in full rescue of 14/16 strains and partial rescue of the remaining 2 strains metG and tyrS (low-background isolates of metG and tyrS were found with additional screening). Escape frequency was calculated by dividing the number of CFU on the restrictive condition (A) by the number of CFU on the permissive condition (D). 12/16 strains had escape frequencies below the limit of detection in the restrictive condition. While dxr and the rpsB isolate shown here exhibit background growth, subsequent screening identified rpsB isolates with strict theophylline dependence (used for quantification in Figure 3). dxr remained conditional in all isolates.

Quantitative characterization of escape frequency revealed a range of performance across the 18 functional strains. Escape frequencies varied from approximately 1 × 10^−5^ to 1 × 10^−6^ (Fig 3) depending on the target gene, which is of comparable single-allele performance to previously described synthetic auxotrophy strategies [5, 30, 31].

**Figure 3:**
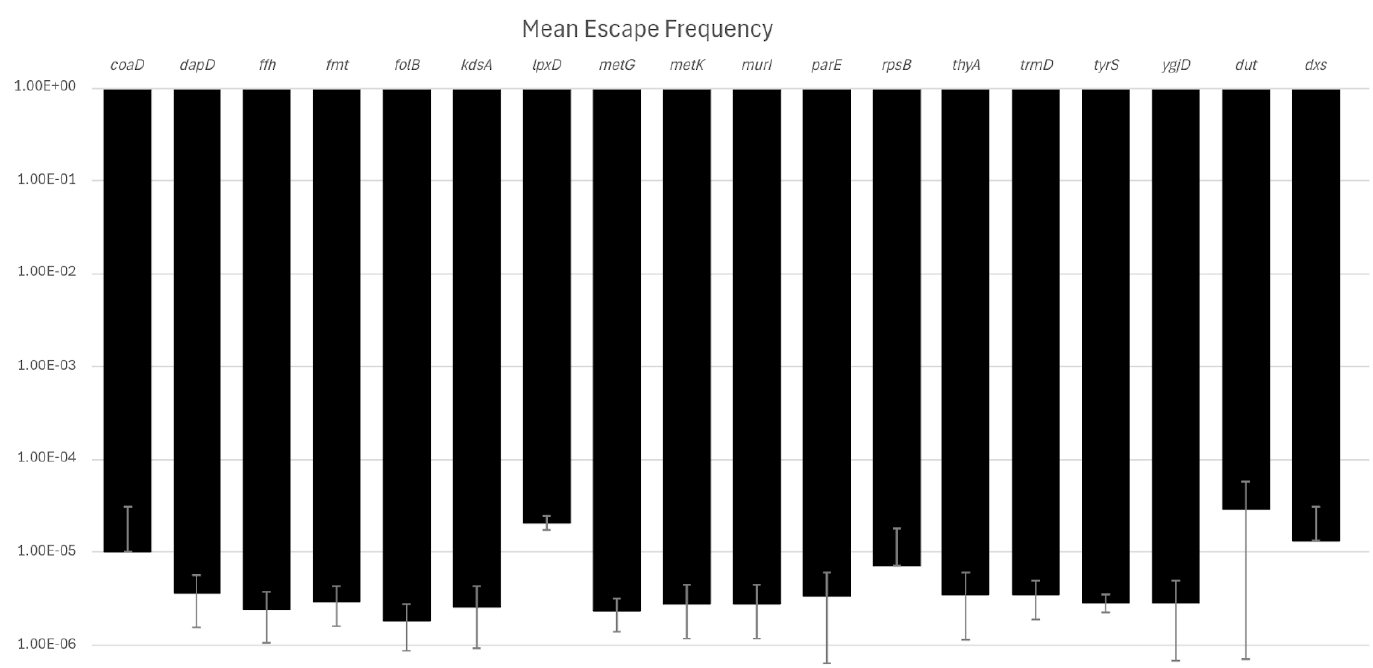
Quantification of Escape Frequencies for Theophylline-Dependent Strains. This bar chart plots the mean escape frequency for the 18 successful synthetic auxotroph strains (dxr is not included as no isolates that were lethal in the restrictive condition were identified). Escape frequency was determined by dividing the number of colonies that grew on restrictive medium (lacking theophylline) by the total number of viable cells plated on permissive medium. The observed mean escape frequencies range from 1.8 × 10-6 (folB) to 2.9 × 10-5 (dut). Most measurements were at the limit of detection (i.e. actual escape frequency may be lower). Each bar represents the mean of 2 to 15 independent biological replicates.

## RESULTS

### Template Design and Strain Construction Methodology

We designed 29 chromosomal knock-in constructs, each targeting a different *E. coli* essential gene, to test whether translational control via a theophylline-responsive riboswitch could be broadly applied to essential genes (Table 1). Targets were chosen from the published Keio collection of *E. coli* essential genes across diverse cellular processes, including DNA replication, translation, cell wall biosynthesis, cofactor metabolism, and lipid synthesis. This approach allowed us to assess whether certain classes of essential genes were more compatible with riboswitch gating of translation.

Each construct consisted of a kanamycin-riboswitch cassette amplified from a plasmid carrying a constitutive promoter (BBa_J72073) driving *kanR* followed by the theophylline-responsive riboswitch riboA (Fig 1A). The cassette was used as a template to create a DNA fragment flanked by essential gene-specific homology arms corresponding to the genomic regions immediately upstream and downstream of each target gene’s start codon (Fig 1D).

Linear PCR fragments were integrated into the chromosome of *E. coli* MC1061 carrying pKD46 using *λ*-Red–mediated recombination (Fig 1E, 1F), with selection on LB agar containing 50 µg/mL kanamycin and 1 mM theophylline [27, 28]. Correct insertions were verified by colony PCR using primers flanking the integration site and confirmed by Sanger sequencing.

Out of the 29 targeted loci, 19 knock-ins were successfully constructed, sequence-verified, and showed clear ligand-dependent phenotypes (18 strict live-die and one high-background conditional; Table 1). Integration efficiency varied substantially between targets: in many cases, transformations yielded hundreds of kanamycin-resistant colonies, whereas in others only a few colonies were recovered. Low-yield of colonies frequently corresponded to integrations that later proved non-conditional, suggesting that locus was particularly sensitive to modification. Several targets produced no colonies even under permissive conditions, implying that riboswitch cassette insertion at those loci disrupted essential gene function and was lethal.

### Phenotypic Outcomes and Classification of Riboswitch-Gated Strains

We evaluated the phenotype of each of the 19 sequence-verified knock-in strains by comparing colony growth under permissive (1 mM theophylline) and restrictive (no theophylline) conditions (Table 1, Fig 2). Eighteen strains exhibited a clear theophylline-dependent viability phenotype, forming normal colonies on LB agar supplemented with theophylline but showing no detectable growth in its absence. These strains represent fully functional theophylline-dependent synthetic auxotrophs, demonstrating that riboswitch-mediated translational control can be successfully imposed across a wide range of essential gene functions. The successful targets included genes involved in cell wall biosynthesis (*murI, lpxD, kdsA*), translation (*tyrS, trmD, rpsB, ffh*), cofactor metabolism (*folB, metK, coaD* ), and DNA replication or maintenance (*parE, dut, thyA*), among others.

In some cases, we observed ligand-modulated growth characterized by reduced, but not abolished, growth in the absence of theophylline. This intermediate phenotype of *dxr* and *rpsB* suggests partial translational repression or residual basal expression sufficient for slow growth (Fig 2A). However, with additional screening of *rpsB*, isolates with low background were easily identified.

The remaining 10 targets yielded non-conditional phenotypes. In cases where knockin efficiencies were high, colonies that grew equivalently on plates with or without theophylline might indicate that the riboswitch failed to effectively gate translation of the essential gene. In cases where knockin efficiencies were low, colonies that grew equivalently on plates with or without theophylline might indicate sensitive loci for which canonical insertions were not viable in the permissive condition.

Variation in phenotypic outcomes across targets likely reflects a combination of gene-specific expression requirements, 5^*′*^ UTR structural context, and mRNA folding constraints imposed by the riboswitch insertion. In essential genes expressed at very high or very low basal levels, the fixed architecture of the riboswitch-kanamycin cassette may have been suboptimal for maintaining the precise expression level or kinetics required for viability. Conversely, failed insertions may have resulted from context incompatibilities, such as disruption of overlapping upstream coding sequences, essential ribosome-binding sites, or promoter elements not captured by the standardized design. Overall, these results demonstrate a success rate of 18 of 29 total targets, supporting the general applicability of riboswitch-based translational control for engineering synthetic auxotrophy across diverse essential gene classes.

## DISCUSSION

### Performance Benchmarking and Contextualization

This work established a new form of synthetic auxotrophy for genetically encoded biocontainment in *E. coli*, based on ligand-dependent translational gating of essential gene products via riboswitches. The platform was designed for speed, modularity, and accessibility, requiring only standard tools like a thermocycler, petri dishes, and the capacity for lambda Red recombineering [27, 28]. Using this modular toolkit, we created theophylline-dependent strains for 18 out of 29 targeted essential genes, demonstrating robust chemical control over viability. These synthetic auxotrophs exhibit strong single-allele performance, with measured escape frequencies of 1 × 10^−5^ to 1 × 10^−6^, a level comparable to other synthetic auxotrophy strategies [5, 30, 31].

### Limitations and Future Directions

This system has several limitations that define its optimal use cases and suggest avenues for future work. First, a single-gene auxotroph with an escape frequency of approximately 1 × 10^−6^, our limit of detection, may not meet the strictest biosafety thresholds for environmental deployment [30]. The true escape frequency is likely lower, but the obvious solution is to stack multiple, orthogonal alleles. As demonstrated in previous works, this approach produces multiplicative decreases in escape frequency, with a two-gene stack predicted to reach a frequency of 1 × 10^−11^ [11, 12, 14].

A second limitation is the toolkit’s reliance on theophylline. While riboswitches offer exceeding modularity and performance [19], the portfolio of ligands used to control riboswitches described in literature is modest compared to protein-ligand interactions.

This platform is designed to lower the barrier for experimenting with genetically encoded biocontainment technologies. While theophylline may not be an ideal compound for all environmental deployment scenarios, this toolkit provides an effective “breadboard” for prototyping synthetic auxotroph biocontainment strategies in the laboratory. By prioritizing simplicity and a low barrier to experimentation, it serves to expand the study of genetically encoded biocontainment, enabling more researchers to rapidly build and test novel concepts.

The modularity of the system also presents clear paths for improving performance. While this study focused only on the riboA riboswitch, future work might increase the number of translationally-gated synthetic auxotrophies by testing different riboswitch variants. Performing basic promoter tuning or incorporating entirely different, well-characterized riboswitches provide many exciting avenues of future investigation [26].

The direct link between riboswitch function and cell viability also provides a powerful selection system to study and engineer nucleic acid-ligand interactions by screening large libraries to evolve riboswitches with novel ligand specificities or an improved dynamic range [22].

As a foundational layer, this system can be integrated into more complex, multi-layered biocontainment strategies to meet diverse safety and containment goals. Escape frequency can be multiplicatively reduced by gating two or more essential genes with independent riboswitches. For multi-modal containment, this translational control can be combined with orthogonal systems, such as post-translational control via SLiDE [14–16] or active kill switches [5, 6, 16]. A powerful dual-control strategy involves using the same riboswitch to gate both an essential gene and an engineered payload circuit, directly linking the circuit’s function to cell viability, addressing concerns related to bio-piracy.

## Conclusion

In conclusion, this work establishes translational initiation control as a viable and effective mechanism for engineering synthetic auxotrophy. We describe a modular toolkit that lowers the barrier to biocontainment by enabling rapid, low-cost implementation with standard molecular biology techniques. The platform is presented with explicit trade-offs, offering a solution that prioritizes ease of implementation and rapid iteration over the bespoke tunability of more complex and resource-intensive systems. This approach expands the mechanistic diversity of developing genetically encoded biocontainment and provides a new engineering option for synthetic biologists.

## MATERIALS AND METHODS

### Bacterial Strains and Growth Conditions

All experiments were performed in *Escherichia coli* strain MC1061 carrying the temperature-sensitive *λ*-Red recombination plasmid pKD46 (carbenicillin resistance). Strains were maintained as 10% (v/v) DMSO stocks at -80°C and streaked onto LB agar (Genesee Scientific, Cat. #69-196) before use. Unless otherwise indicated, cultures were grown in 2xYT broth (Genesee Scientific, Cat. #69-103) or on LB agar at 37 °C with shaking at 250 rpm.

Theophylline monohydrate (Thomas Scientific, Cat. #C950H07) was prepared as a 20 mM stock solution in sterile deionized water and filter-sterilized. For permissive growth conditions, theophylline was added to media at a final concentration of 1 mM. Kanamycin monosulfate (GoldBio, Cat. #K-120-5) was used at 50 µg/mL for selection of kanR-containing strains. When required to maintain pKD46, carbenicillin (200 µg/mL) was included.

Cells harboring pKD46 were incubated at 30 °C to preserve the temperature-sensitive plasmid. For *λ*-Red induction, 10 mM L-arabinose was added at mid-log phase (OD_600_ ≈ 0.4). Optical density at 600 nm was measured using a Spectronic 20 Genesys spectrophotometer. All other cultures and phenotypic assays were performed at 37 °C unless otherwise noted.

### Molecular Cloning and Plasmid Construction

The riboswitch–selection cassette used for all genomic integrations was derived from plasmid bgg241, which contains a kanamycin resistance marker driven by the constitutive promoter BBa_J72073 (Registry of Standard Biological Parts, https://parts.igem.org/Part:BBa_J72073). A theophylline-responsive riboswitch was inserted immediately downstream of the *kanR* coding sequence in bgg241 by Golden Gate assembly. The theophylline dependent riboswitch sequence was obtained from Topp et al [23].

To generate gene-specific integration fragments, PCR amplification was performed using Q5 High-Fidelity DNA Polymerase (New England Biolabs) and primers containing approximately 50 bp homology arms corresponding to the genomic regions immediately upstream and downstream of each target gene’s start codon. The homology arms were encoded directly within the primers, which were synthesized by Integrated DNA Technologies (IDT). PCR products were purified using the Zymo Research DNA Clean & Concentrator Kit and verified by agarose gel electrophoresis before electroporation into *λ*-Red containing *E. coli* cells for chromosomal integration

Primer sequences are listed in S2 Table.

### Lambda Red Recombination for Genomic Integration

Gene-specific kanamycin-riboswitch cassettes were integrated into the *E. coli* chromosome using *λ*-Red mediated homologous recombination. Recombineering was performed in *E. coli* strain MC1061 carrying the temperature-sensitive helper plasmid pKD46, which encodes the *λ*-Red recombination genes..

Cells harboring pKD46 were grown overnight at 30 °C in 2xYT medium containing 200 µg/mL carbenicillin, then subcultured 1:100 into fresh 2xYT supplemented with 10 mM L-arabinose to induce *λ*-Red expression. Cultures were incubated at 30 °C until reaching an optical density of OD_600_ ≈ 0.4, then chilled on ice and washed three times with ice-cold 10% glycerol to prepare electrocompetent cells.

Purified linear PCR fragments containing the riboswitch cassette and gene-specific homology arms were electroporated into 90 µL of electrocompetent cells using a 0.1 cm gap cuvette at 1.6 kV. The total amount of DNA introduced varied depending on yield. Following electroporation, cells were immediately recovered in 2xYT medium supplemented with 1 mM theophylline and incubated for 1 hour at 30 °C with shaking to allow recombination and expression of antibiotic resistance.

Transformants were selected on LB agar plates containing 50 µg/mL kanamycin and 1 mM theophylline, and incubated overnight at 37 °C. Colony PCR was performed to confirm correct integration of the cassette at the targeted locus using primers flanking the recombination site. Verified colonies were streak-purified and stored as 10% DMSO freezer stocks at -80 °C.

### Phenotypic Characterization and Escape Frequency Measurements

To evaluate ligand-dependent growth and quantify escape frequency, single colonies were picked from selective plates and resuspended in phosphate-buffered saline (PBS). The suspensions were serially diluted in 10-fold increments in PBS, and 10 µL of each dilution was spotted onto LB agar plates containing 50 µg/mL kanamycin, either with (1 mM theophylline) or without theophylline.

Plates were incubated overnight at 37 °C, after which the colonies in each spot were counted and recorded as colony-forming units per milliliter (CFU/mL). Escape frequency was calculated as the ratio of CFU/mL observed on restrictive plates (without theophylline) to that on permissive plates (with theophylline) for the same strain.

Each strain was measured using 2-15 independent biological replicates, depending on the number of available transformants. For strains exhibiting no detectable colonies under restrictive conditions, escape frequency was reported as below the limit of detection and a maximum possible CFU/mL value corresponding to the detection limit (e.g., 99 CFU/mL) was reported for transparency in calculations.

## Acknowledgments

We thank John Christopher Anderson for Discussion and Review

## Data Availability Statement

All data underlying the results reported in this manuscript are available within the manuscript and its Supporting Information files. There are no restrictions on data availability.

## Ethics Statement

This study did not involve human participants, vertebrate animals, or regulated invertebrate species. All experiments were performed in vitro using microbial cultures, for which institutional ethics committee approval is not required under applicable institutional and national guidelines. No additional ethics approval was therefore required.

## Supplementary

**S1 Table.**
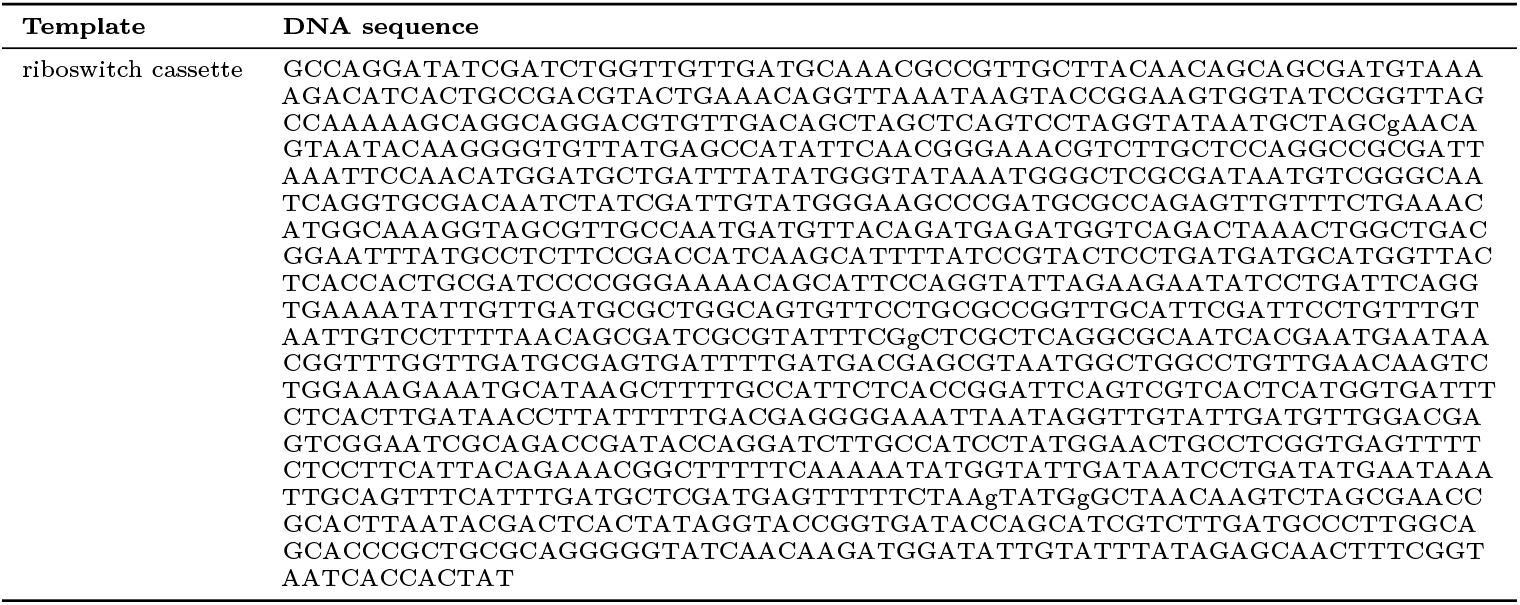
Kanamycin Resistance Marker and riboA cassette template. This table provides the DNA sequence for the kanR-riboA cassette that is used as template for PCR amplification of knock-in ready fragments targeting upstream of desired essential genes.

**S2 Table.**
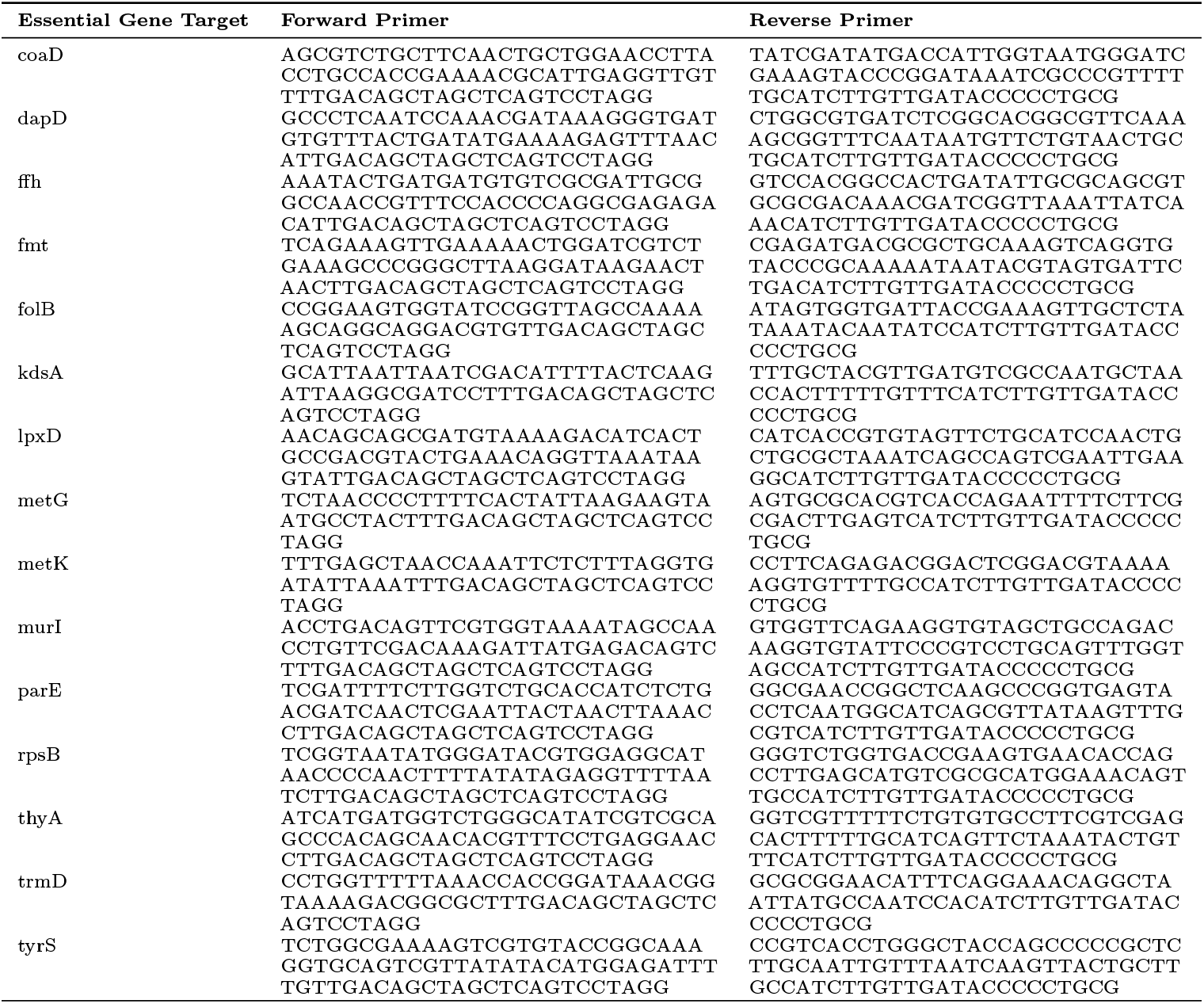
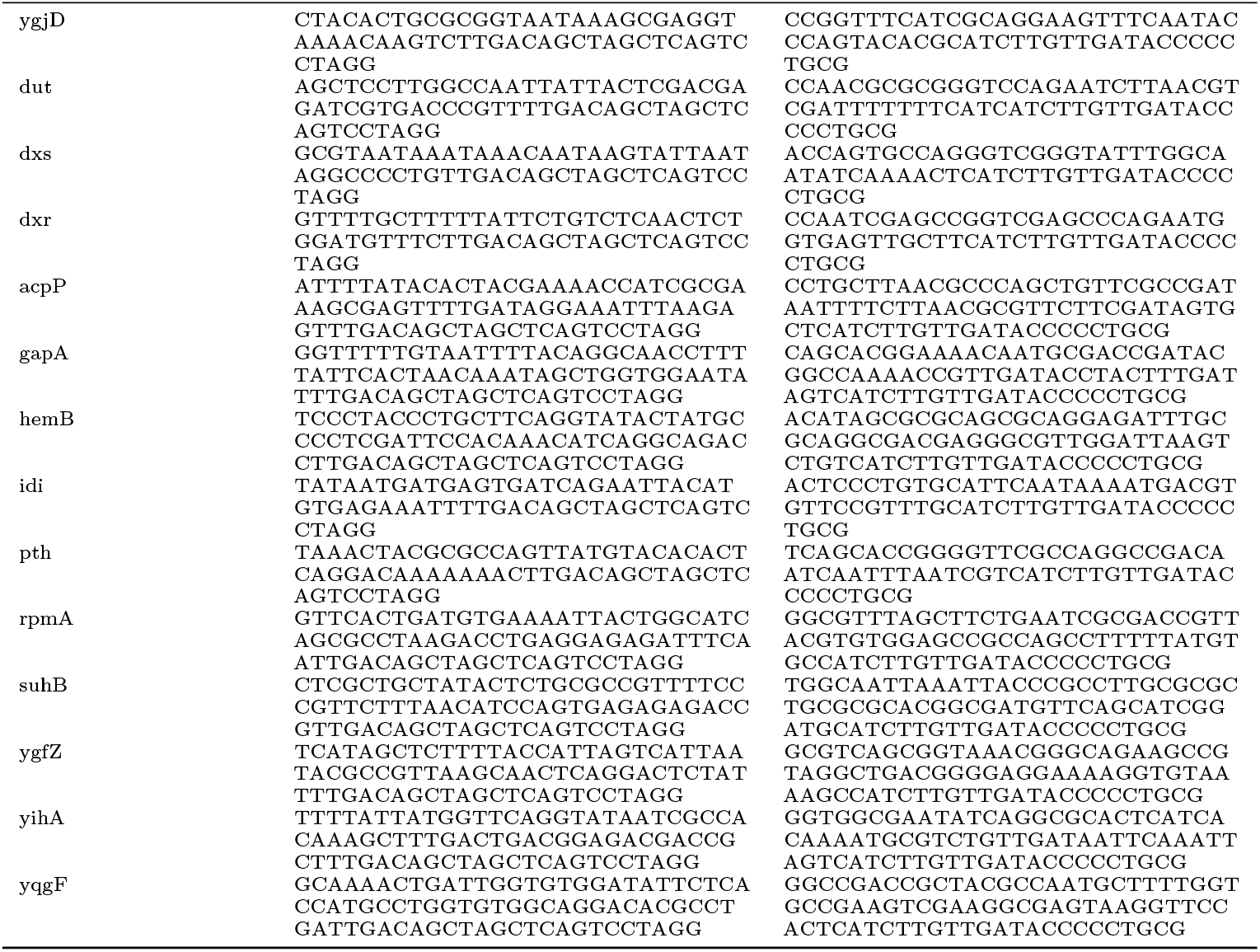
Primers used in this study to create the kanR-riboA knock-in fragments. This table provides DNA primers used to amplify knockin fragments (from template provided in S1 Table). The annealing region targets the kanR-riboA fragment and the 5 prime of each primer provides homology targeting the upstream sequence of 29 essential E. coli genes.

